# Cardiovascular Disease Risk Factor Responses to a Type 2 Diabetes Care Model Including Nutritional Ketosis at One Year: An Open Label, Non-Randomized, Controlled Study

**DOI:** 10.1101/262709

**Authors:** Nasir H. Bhanpuri, Sarah J. Hallberg, Paul T. Williams, Amy L. McKenzie, Kevin D. Ballard, Wayne W. Campbell, James P. McCarter, Stephen D. Phinney, Jeff S. Volek

## Abstract

**Background**

Cardiovascular disease (CVD) is a leading cause of death among adults with type 2 diabetes mellitus (T2D). We recently reported that glycemic control in patients with T2D can be significantly improved through a continuous care intervention (CCI) including nutritional ketosis. The purpose of this study was to examine CVD risk factors in this cohort.

**Methods**

We investigated CVD risk factors in patients with T2D who participated in a one year open label, non-randomized, controlled study. The CCI group (n = 262) received treatment from a health coach and medical provider. A usual care (UC) group (n = 87) was independently recruited to track customary T2D progression. Circulating biomarkers of cholesterol metabolism and inflammation, blood pressure (BP), carotid intima media thickness (cIMT), multi-factorial risk scores and medication use were examined.

**Results**

The CCI group consisted of 262 patients (baseline mean(SD): age 54(8) y, BMI 40.4(8.8) kg/m^2^). Intention-to-treat analysis (% change) revealed the following at 1-year with P values < 0.0019 indicating statistical significance after adjustment for multiple comparisons: total LDL-particles (LDL-P) (−4.9%, P=0.02), small LDL-P (−20.8%, P=1.2×10^−12^), LDL-P size (+1.1%, P=6.0×10^−10^), ApoB (−1.6%, P=0.37), ApoA1 (+9.8%, P<10^−16^), ApoB/ApoA1 ratio (−9.5%, P=1.9×10^−7^), triglyceride/HDL-C ratio (−29.1%, P<10^−16^), large VLDL-P (−38.9%, P=4.2×10^−15^), and LDL-C (+9.9%, P=4.9×10^−5^). Additional effects were reductions in blood pressure, high sensitivity C-reactive protein, and white blood cell count (all P<1×10^−7^) while cIMT was unchanged. The 10-year atherosclerotic cardiovascular disease (ASCVD) risk score decreased −11.9% (P=4.9×10^−5^). Antihypertensive medication use was discontinued in 11.4 % of CCI participants (P=5.3×10^−5^). The UC group of 87 patients (baseline mean(SD): age 52(10)y, BMI 36.7(7.2) kg/m^2^) showed no significant changes. After adjusting for baseline differences when comparing CCI and UC groups, significant improvements for the CCI group included small LDL-P, ApoA1, triglyceride/HDL-C ratio, HDL-C, hs-CRP, and ASCVD score. The CCI group showed a greater rise in LDL-C.

**Conclusions**

A continuous care treatment including nutritional ketosis in patients with T2D improved most biomarkers of CVD risk after one year. The increase in LDL-cholesterol appeared limited to the large LDL subfraction. LDL particle size increased, total LDL-P and ApoB were unchanged, and inflammation and blood pressure decreased.

**Trial registration**

Clinicaltrials.gov: NCT02519309. Registered 10 August 2015

## Background

Despite advances in the prevention and treatment of cardiovascular disease (CVD), it remains the leading cause of death in adults across the world [1]. Specifically, among those with type 2 diabetes (T2D) in the U.S., CVD accounts for 44% of mortality [2]. T2D rates have doubled over the past twenty years [3] and CVD risk increases two to four-fold with a diagnosis of T2D [4], warranting the identification of novel interventions to combat T2D. Intensive lifestyle interventions with dietary carbohydrate restriction [5-8], including the recently described continuous remote care model, which helps patients with T2D sustain nutritional ketosis [9,10], have demonstrated improved glycemic control concurrent with medication reduction. However, the long-term sustainability and impact of these interventions on CVD risk and lipid profiles remains a subject of debate [11,12].

Atherogenic dyslipidemia, a known risk factor for CVD [13], is highly prevalent in patients with T2D [14] and tightly linked to high-carbohydrate diets[15]. The condition is characterized by increased triglycerides, decreased high-density lipoprotein cholesterol concentration (HDL-C) and increased small low-density lipoprotein particle number (small LDL-P). Evidence suggests that increased very low-density lipoprotein particle number (VLDL-P), and in particular large VLDL-P, may be one of the key underlying abnormalities in atherogenic dyslipidemia [14,16-18]. Elevated concentrations of small LDL are often associated with increased total LDL particle number (LDL-P) and ApoB [19,20]. Particularly in patients with insulin resistance and T2D, elevated LDL-P and ApoB may exist even with normal to low LDL-C values [19,21,22]. Reliance on LDL-C for risk assessment in T2D patients may miss the impact of atherogenic dyslipidemia and elevated LDL-P. It has been proposed that LDL-P or ApoB may be superior to LDL-C as a predictor of CVD [22-25].

Previous studies of carbohydrate restriction of up to 1-year found a consistent decrease in triglycerides and increase in HDL-C, while LDL-C slightly increased or decreased [15,26-28]. Although LDL-C is a risk factor for CVD, low LDL-C may belie elevations in small LDL, LDL-P or ApoB. Conversely, increased LDL-C with a low carbohydrate diet may primarily reflect the large LDL subfraction and may not increase CVD risk if total LDL-P or ApoB concentrations are unchanged or decline.

Inflammation, as assessed by elevated high-sensitivity C-reactive protein (hsCRP) or white blood cell count (WBC) [29-32], is an independent CVD risk factor and is involved in all stages of atherogenesis [33]. Inflammation is often observed in T2D concurrent with atherogenic dyslipidemia [34] and represents an additional CVD risk even in individuals with low to normal LDL-C [35,36]. Hypertension is an additive risk factor in this patient population. Tighter blood pressure control has been associated with reduction in the risk of deaths related to diabetes. This included decreased CVD, stroke and microvascular complications [37].

For this open label, non-randomized, controlled, before-and-after study, we investigated the effects of a continuous care intervention (CCI) on CVD risk factors. The CCI included individualized digital support with telemedicine, health coaching, education in nutritional ketosis, biometric feedback, and an online peer-support community. Given the multi-faceted pathophysiology of CVD, we assessed the one-year responses in several biomarkers related to cholesterol and lipoprotein metabolism, blood pressure, and inflammation, as well as carotid intima media thickness (cIMT) and medication use. Some results were previously reported in relation to glycemic control [10] and are presented here as they pertain to CVD risk.

## Methods

### Intervention

As previously described [9,10], we utilized a prospective, longitudinal study design with a cohort of patients with T2D from the greater Lafayette, Indiana, USA, region who self-selected to participate in the CCI (Clinicaltrials.gov Identifier NCT02519309). Patients in the CCI were provided access to a web-based software application (app) for biomarker reporting and monitoring including body weight, blood glucose and blood betahydroxybutyrate (BHB; a marker of ketosis). The remote care team consisted of a health coach and physician or nurse practitioner who provided nutritional advice and medication management, respectively. Patients were guided by individualized nutrition recommendations to achieve and sustain nutritional ketosis. CCI participants self-selected to receive education via either an onsite group setting (CCI-onsite) or via the app (CCI-web). Social support was provided via an online peer community. This study was approved by the Franciscan Health Lafayette Institutional Review Board, and participants provided written informed consent.

Glycemic control medication management was previously described [9,10]. For participants with a history of hypertension, a home automatic sphygmomanometer was supplied. Participants reported their home readings in the app daily to weekly depending on recent control and instruction from the supervising physician. Antihypertensive prescriptions were adjusted based on home readings and reported symptoms. Health coaches responded to patient app reported readings of systolic blood pressure less than 110 mmHg with specific questions about symptoms of hypotension. Following resolution of hypertension, diuretics were the first antihypertensive medications to be discontinued, followed by beta blockers, unless the participant had a history of coronary artery disease. Angiotensin-converting-enzyme inhibitors (ACEs) and angiotensin II receptor blockers (ARBs) were generally continued due to known renal protection with diabetes [38,39]. Statin medications were adjusted when appropriate to maintain a goal of LDL-P under 1000 nmol·L^-1^ or participant preference after full risk-benefit discussion.

To track T2D progression in the same geography and health system as the CCI, an independent cohort of patients with T2D who received usual care (UC) were recruited. These patients were referred to registered dietitians providing dietary advice according to American Diabetes Association guidelines [40].

### Outcome Measures

Anthropometrics and vital signs for the CCI group were obtained at baseline, 70 days, and one year. A stadiometer was used to assess height and used in the calculation of body mass index. A calibrated scale in the clinic measured weight to the nearest 0.1 lb (Model 750, Detecto; Webb City, MO, USA) and values were converted to kg. Participants were provided a cellular-connected home scale for daily weight. Blood pressure was obtained manually by trained staff after participants rested in a seated position for five minutes. Adverse events were reported and reviewed by the Principal Investigator and the Institutional Review Board.

Fasting blood draws for the CCI group were collected at baseline, 70 days, and one-year follow-up (ranging from 11 - 15 months). Blood analytes were determined via standard procedures at a Clinical Laboratory Improvement Amendment (CLIA) accredited laboratory on the day of sample collection or from stored serum. Serum aliquots were stored at −80°C and thawed for determination of ApoB, ApoA1, total cholesterol, triglycerides, and direct HDL-C concentrations by FDA approved methods (Cobas c501, Roche Diagnostics; Indianapolis, IN, USA). LDL was calculated using the Friedewald equation [41]. Lipid subfractions were quantified using clinical NMR LipoProfile^®^ (LabCorp, Burlington NC, USA; [42]). The LipoProfile3 algorithm used in the present investigation was used previously to relate lipid subfractions to CVD risk [35,43,44]. The NMR-derived lipoprotein insulin resistance score (LP-IR) is proposed to be associated with the homeostasis model assessment of insulin resistance (HOMA-IR) and glucose disposal rate (GDR) [45]. The multifactorial 10-year atherosclerotic cardiovascular disease (ASCVD) risk score was also computed [46].

Anthropometrics, vital signs and fasting blood draws for the UC group were obtained at baseline and one year as described above using the same clinical facilities and laboratory and data collection methods. Home biometrics for the UC group were not tracked and 70-day outcomes were not measured.

Carotid ultrasonography for cIMT measure was performed at baseline and one year in CCI and UC groups to characterize atherosclerotic risk. Ultrasound technicians were trained according to protocols that were previously tested and used to assess subclinical atherosclerosis [47,48]. The right and left common carotid arteries were imaged one cm distal to the carotid bulb using a L12-3 multi-frequency linear-array transducer attached to a high-resolution ultrasound system (Phillips EPIQ 5, Amsterdam, Netherlands). Longitudinal images were captured in three imaging planes: anterior, lateral, and posterior. Digital images were analyzed using edge-detection software (Carotid Analyzer for Research; Medical Imaging Application, Coralville, IA) to trace the lumen-intima and intima-medial boundaries of the artery. Analyses were performed by the same blinded investigator to obtain right and left mean arterial diameter and mean cIMT.

### Statistics

JMP software (version 5.1, SAS Institute; Cary, SC, USA) was used for all statistical analyses except multiple imputation. Multiple imputation, conducted with Stata software (version 11, StataCorp; College Station, TX, USA), was used to estimate means and standard errors describing the variability between imputations. Seven hundred imputations from multivariate normal regression were run to estimate the missing values (4% missing at baseline and 22% missing at one year). Two-sample t-tests were used to test for significance of the differences in baseline biomarker values between groups. Two-sample t-tests were also used to test for differences between one-year changes between groups. Paired t-tests and analysis of covariance (ANCOVA) when adjusted for baseline covariates (sex, age, baseline BMI, and African-American race) were used to test for significance of within-group changes. To reduce skewness before testing for significance, triglyceride, triglyceride/HDL-C ratio and hsCRP were first log-transformed, however aside from P values, the tables present results from the untransformed data. Percent change in a given biomarker was calculated as the mean difference value divided by the mean baseline value. The standard error of percent change of a given biomarker was calculated as the standard error of the change divided by the mean baseline value. Significant changes in proportions (e.g. medication use) were tested using McNemar’s test with continuity correction in completers, and linear regression of the changes in the dichotomous states when missing outcome data were imputed.

Throughout the manuscript, standard deviations are presented within parentheses and standard errors are presented following “±” symbol. Nominal significance levels (P) are presented in the tables; however, a significance level of P<0.0019 ensures simultaneous significance at P<0.05 for a Bonferroni adjustment for the 26 variables examined. Unless otherwise noted, results presented are intention-to-treat analyses (all starters) with missing values estimated by imputation. Some results are designated as completer analyses (excluding participants who withdrew or lacked biomarkers at one year).

## Results

### Baseline characteristics of participants

The baseline characteristics of the 262 T2D participants who began the CCI are shown in Table 1. There were no significant differences in baseline characteristics between groups self-selecting web-based (CCI-web) and onsite education (CCI-onsite) (Supplementary Table 1) nor were there significant differences in biomarker changes at one year between the groups (Supplementary Table 2). Therefore, results for the two groups were combined for further analyses.

**Table 1.**
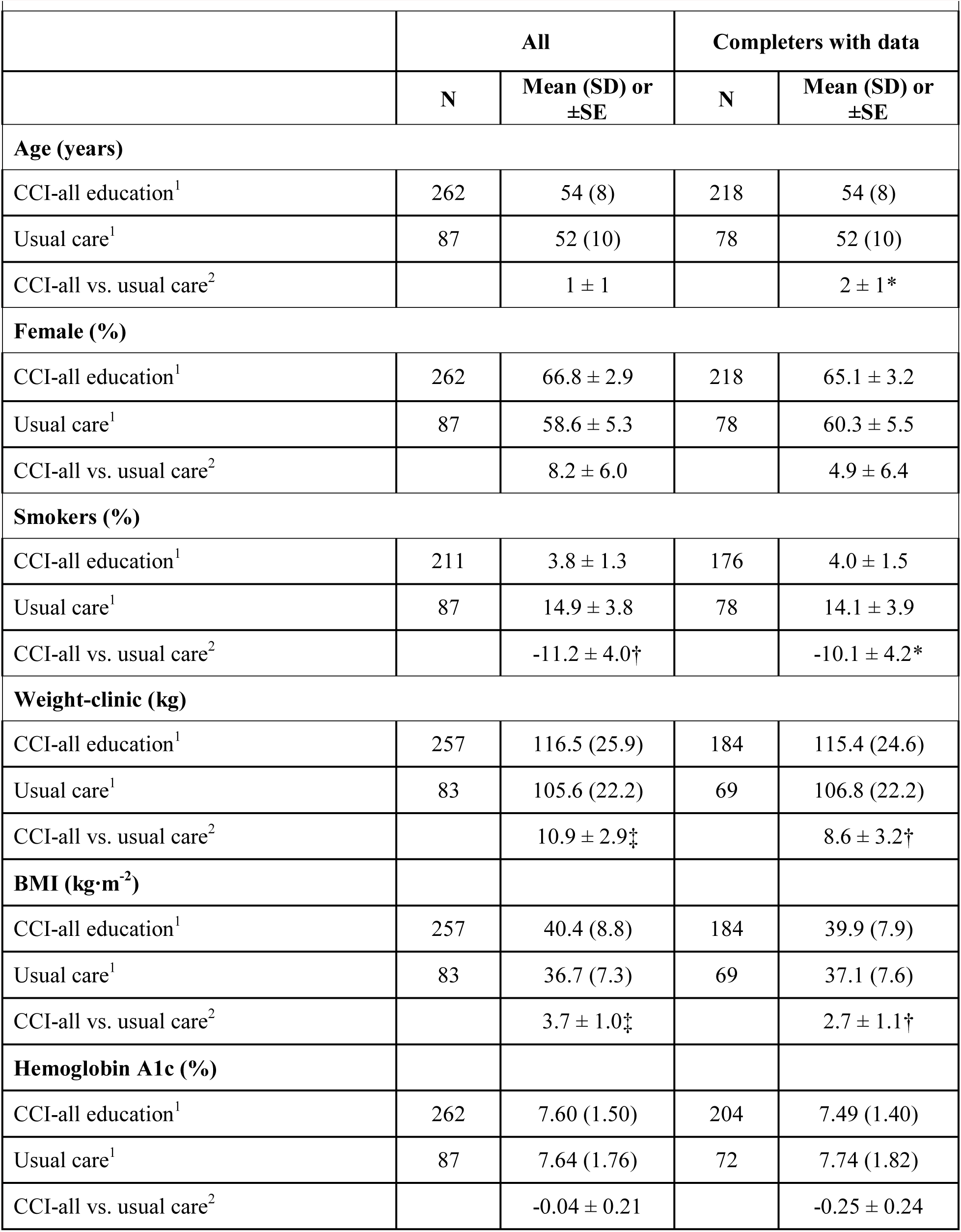
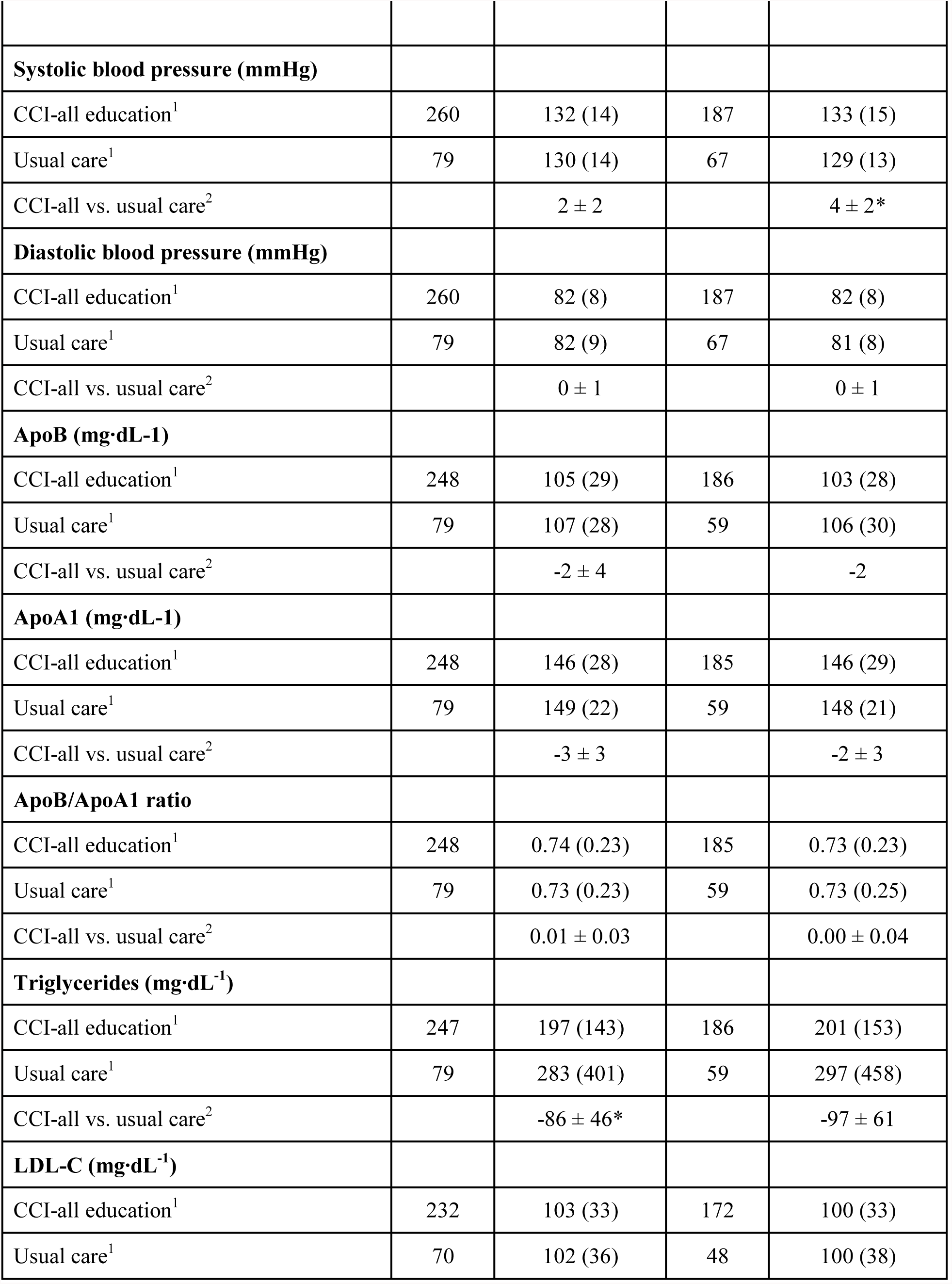
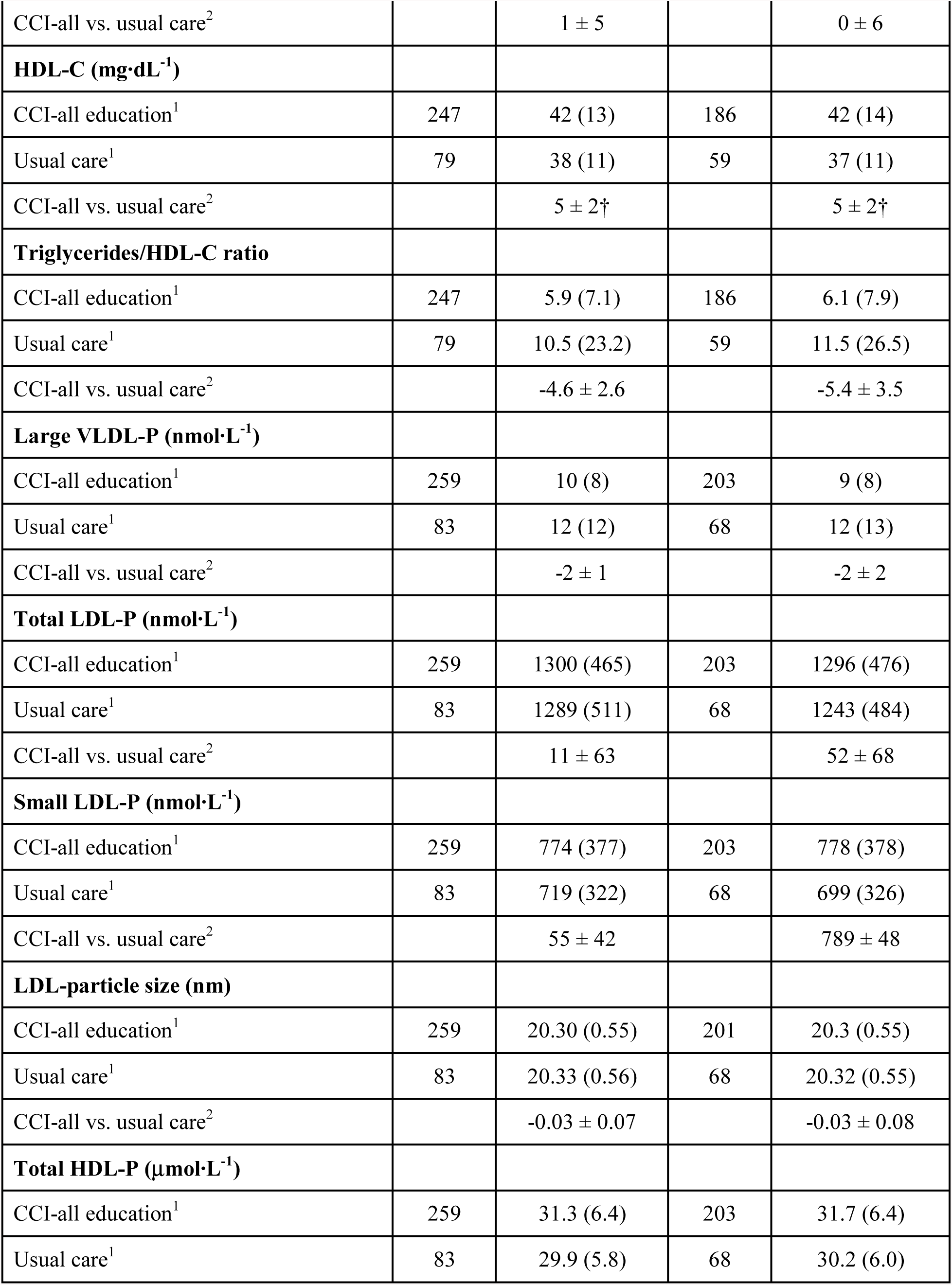
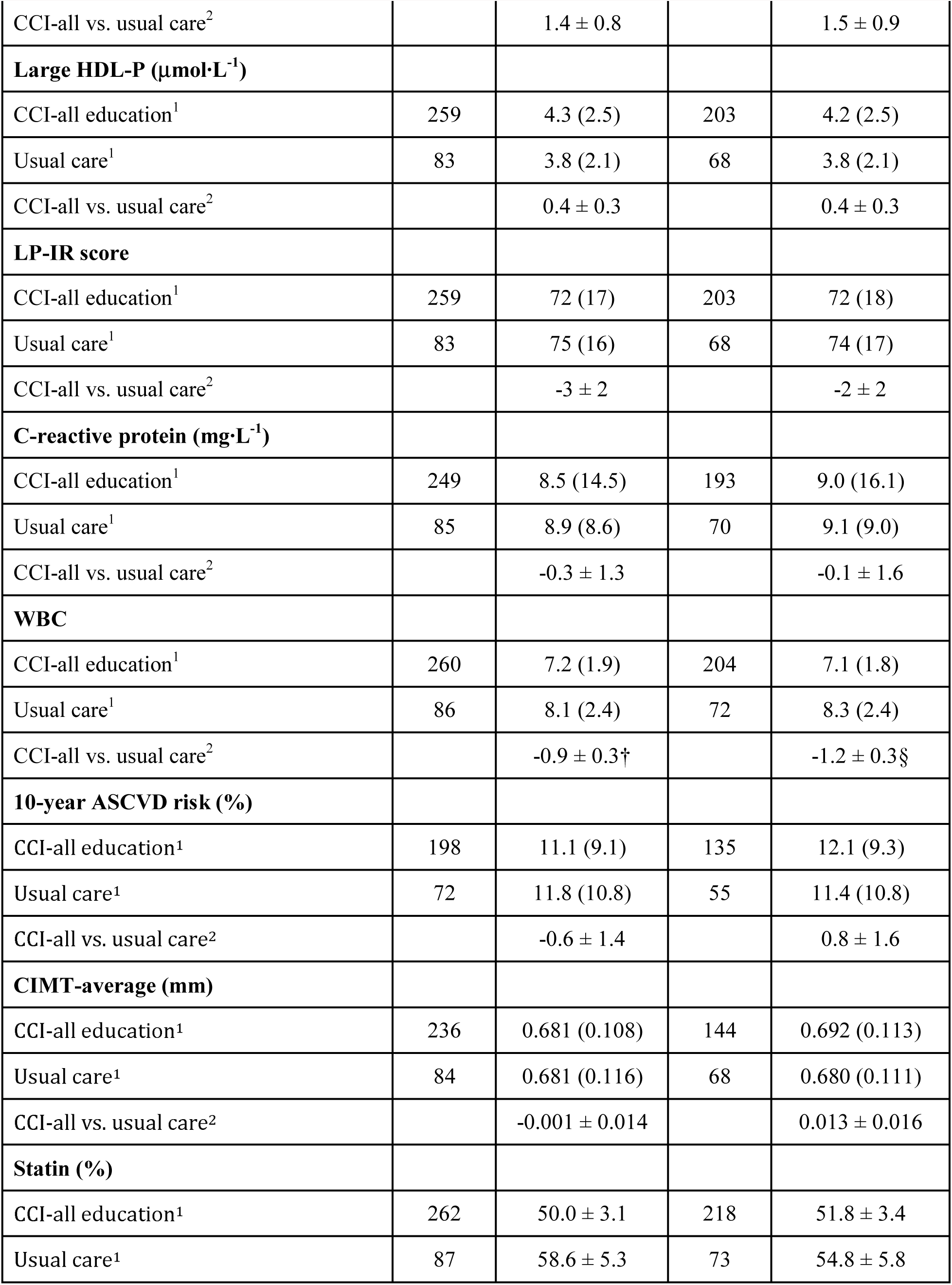
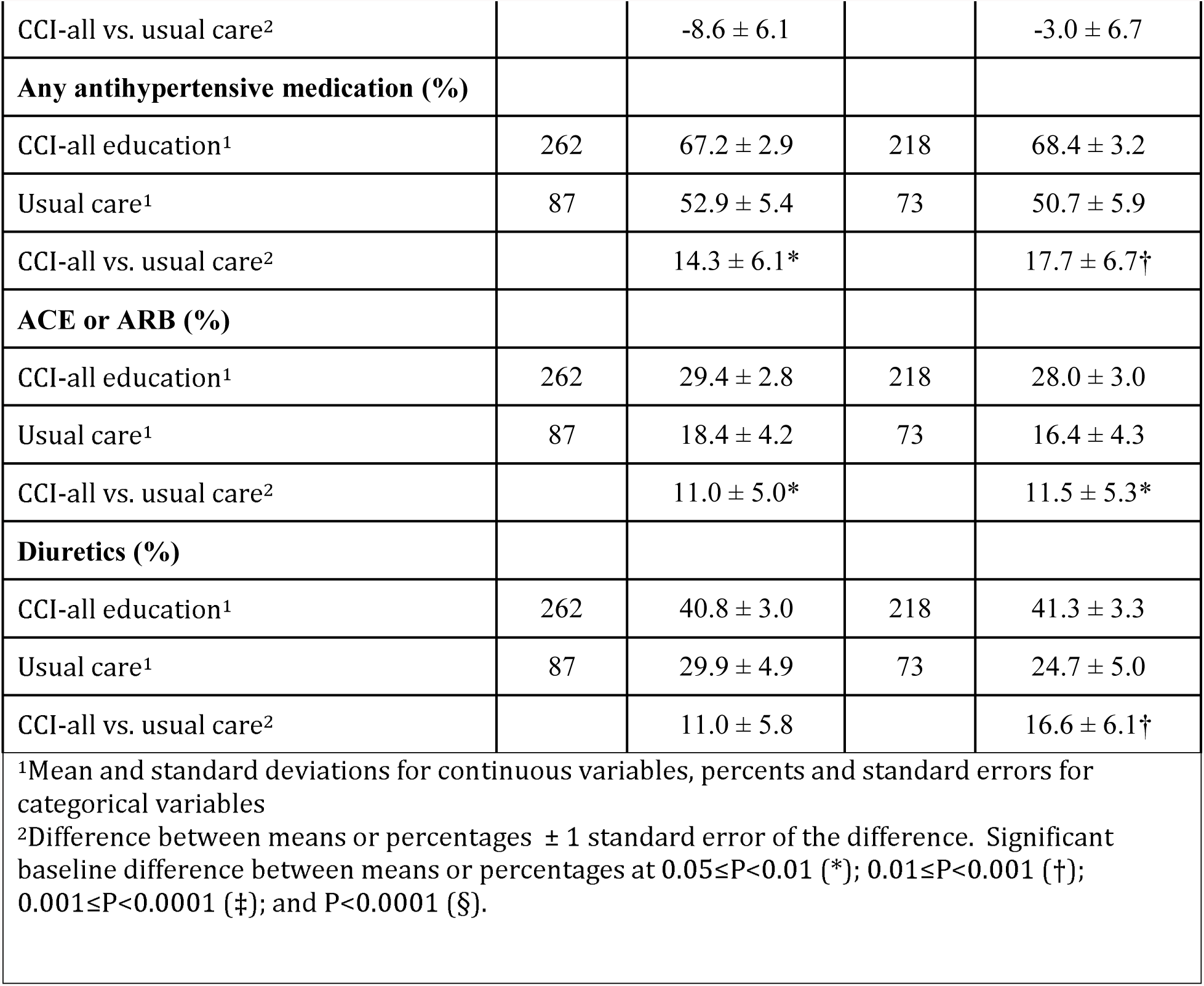
Baseline characteristics Continuous Care Intervention (CCI) and Usual Care (UC) groups.

The baseline characteristics of participants with measurements at both baseline and one year were not significantly different from dropouts and participants with missing data after correcting for multiple comparisons (Supplementary Table 1).

An independently recruited UC group of 87 T2D participants, which provided an observational comparison group from the same geography and health system, showed no significant differences from the CCI group in baseline characteristics except mean body weight and BMI were higher in the CCI versus the UC group (Table 1, P<0.001).

### Changes in biomarkers of CVD risk at one year

Two-hundred eighteen (83%) participants remained enrolled in the CCI group at one year. One-year changes in CVD biomarkers are detailed in Table 2 and percent changes from baseline are shown in Figure 1. The within-CCI group changes in the following lipids and lipoproteins were all statistically significant after adjusting for multiple comparisons (P<0.0019), reported here as mean percent difference from baseline: ApoA1 (+9.9%), ApoB/ApoA1 ratio (−9.5%), triglycerides (−24.4%), LDL-C (+9.6%), HDL-C (+18.1%), triglyceride/HDL-C ratio (−29.1%), large VLDL-P (−38.9%), small LDL-P (−20.8%), LDL-particle size (+1.1%), total HDL-P (+4.9%), and large HDL-P (+23.5%). There were no significant changes after adjusting for multiple comparisons in total LDL-P (−4.9%, p=0.02) or ApoB (−1.6%, p=0.37).

**Table 2.** One-year Biomarker Change Continuous Care Intervention compared to Usual Care groups.

|  | Completers |  |  |  |  |  | All starters<br>(Dropouts imputed)<sup>4</sup> |  |  |
| --- | --- | --- | --- | --- | --- | --- | --- | --- | --- |
|  | N |  | Unadjusted |  | Adjusted for baseline<sup>3</sup> |  | Unadjusted |  |  |
| | | One Year<br>Mean ±SE | Difference<br>(SD) or<br>±SE | Signif-<br>icance<sup>5</sup> | Difference<br>±SE | Signif-<br>icance<sup>5</sup> | One Year<br>Mean ±SE | Difference<br>±SE | Signif-<br>icance<sup>5</sup> |
| **ΔWeight-clinic (kg)** |  |  |  |  |  |  |  |  |  |
| CCI-all education<sup>1</sup> | 184 | 101.2 ± 1.6 | -14.2<br>(10.3) | <10<sup>-16</sup> | -13.8 ± 0.6 | <10<sup>-16</sup> | 102.7 ± 1.5 | -13.8 ± 0.7 | <10<sup>-16</sup> |
| Usual care<sup>1</sup> | 69 | 106.8 ± 2.7 | 0.04 (5.9) | 0.95 | -1.1 ± 1.1 | 0.29 | 107.3 ± 2.6 | -0.2 ± 0.8 | 0.85 |
| CCI-all vs. usual care<sup>2</sup> | | | -14.3 ± 1.0 | <10<sup>-16</sup> | -12.7 ± 1.3 | <10<sup>-16</sup> | | -13.7 ± 1.1 | <10<sup>-16</sup> |
| **ΔHemoglobin A1c (%)** |  |  |  |  |  |  |  |  |  |
| CCI-all education<sup>1</sup> | 204 | 6.20 ± 0.07 | -1.29<br>(1.32) | <10<sup>-16</sup> | -1.32 ±<br>0.09 | <10<sup>-16</sup> | 6.29 ± 0.07 | -1.30 ±<br>0.09 | <10<sup>-16</sup> |
| Usual care<sup>1</sup> | 72 | 7.94 ± 0.22 | 0.20 (1.35) | 0.21 | 0.22 ± 0.16 | 0.17 | 7.84 ± 0.19 | 0.20 ± 0.15 | 0.18 |
| CCI-all vs. usual care<sup>2</sup> | | | -1.49 ±<br>0.18 | 4.4x10<sup>-16</sup> | -1.54 ±<br>0.19 | 4.4x10<sup>-16</sup> | | -1.50 ±<br>0.17 | <10<sup>-16</sup> |
| **ΔSystolic blood pressure**<br>**(mmHg)** |  |  |  |  |  |  |  |  |  |
| CCI-all education<sup>1</sup> | 187 | 126 ± 1 | -7 (16) | 1.3x10<sup>-8</sup> | -7 ± 1 | 1.6x10<sup>-7</sup> | 126 ± 1 | -6 ± 1 | 1.3x10<sup>-8</sup> |
| Usual care<sup>1</sup> | 67 | 129 ± 2 | 0 (18) | 0.91 | 0 ± 2 | 0.83 | 129 ± 2 | -1 ± 2 | 0.67 |
| CCI-all vs. usual care<sup>2</sup> | | | -7 ± 2 | 0.005 | -6 ± 3 | 0.02 | | -5 ± 2 | 0.02 |
| **ΔDiastolic blood pressure**<br>**(mmHg)** |  |  |  |  |  |  |  |  |  |
| CCI-all education<sup>1</sup> | 187 | 78 ± 1 | -4 (9) | 1.4x10<sup>-7</sup> | -4 ± 1 | 6.2x10<sup>-7</sup> | 79 ± 1 | -4 ± 1 | 7.2x10<sup>-8</sup> |
| Usual care<sup>1</sup> | 67 | 81 ± 1 | 0 (10) | 0.92 | 0 ± 1 | 0.75 | 81 ± 1 | -1 ± 1 | 0.45 |
| CCI-all vs. usual care<sup>2</sup> |  |  | -3 ± 1 | 0.01 | -3 ± 1 | 0.03 |  | -3 ± 1 | 0.06 |
| **ΔApoB (mg·dL<sup>-1</sup>)** |  |  |  |  |  |  |  |  |  |
| CCI-all education<sup>1</sup> | 186 | 103 ± 2 | -1 (24) | 0.69 | -0 ± 2 | 0.82 | 104 ± 2 | -2 ± 2 | 0.37 |
| Usual care<sup>1</sup> | 59 | 107 ± 5 | 2 (37) | 0.75 | 1 ± 4 | 0.9 | 106 ± 4 | 0 ± 4 | 0.95 |
| CCI-all vs. usual care<sup>2</sup> |  |  | -2 ± 5 | 0.66 | -1 ± 5 | 0.83 |  | -2 ± 5 | 0.67 |
| **ΔApoA1 (mg·dL<sup>-1</sup>)** |  |  |  |  |  |  |  |  |  |
| CCI-all education<sup>1</sup> | 185 | 160 ± 3 | 14 (24) | 8.9x10<sup>-16</sup> | 14 ± 2 | 4.4x10<sup>-16</sup> | 160 ± 2 | 14 ± 2 | <10<sup>-16</sup> |
| Usual care<sup>1</sup> | 59 | 145 ± 3 | -3 (19) | 0.18 | -2 ± 3 | 0.55 | 147 ± 3 | -2 ± 3 | 0.37 |
| CCI-all vs. usual care<sup>2</sup> |  |  | 18 ± 3 | 4.7x10<sup>-9</sup> | 16 ± 4 | 2.2x10<sup>-5</sup> |  | 17 ± 3 | 1.4x10<sup>-7</sup> |
| **ΔApoB/ApoA1** |  |  |  |  |  |  |  |  |  |
| CCI-all education<sup>1</sup> | 185 | 0.67 ± 0.02 | -0.06<br>(0.17) | 1.8x10<sup>-6</sup> | -0.06 ±<br>0.02 | 0.003 | 0.67 ± 0.02 | -0.07 ±<br>0.01 | 1.9x10<sup>-7</sup> |
| Usual care<sup>1</sup> | 59 | 0.76 ± 0.04 | 0.03 (0.29) | 0.42 | 0.02 ± 0.03 | 0.5 | 0.74 ± 0.03 | 0.02 ± 0.03 | 0.58 |
| CCI-all vs. usual care<sup>2</sup> |  |  | -0.09 ±<br>0.04 | 0.02 | -0.08 ±<br>0.03 | 0.02 |  | -0.09 ±<br>0.03 | 0.01 |
| **ΔTriglycerides (mg·dL<sup>-1</sup>)** |  |  |  |  |  |  |  |  |  |
| CCI-all education<sup>1</sup> | 186 | 151 ± 11 | -49 (168) | 5.6x10<sup>-5</sup> | -50 ± 16 | 0.001 | 148 ± 12 | -48 ± 13 | <10<sup>-16</sup> |
| Usual care<sup>1</sup> | 59 | 327 ± 65 | 30 (301) | 0.44 | 31 ± 29 | 0.27 | 305 ± 48 | 28 ± 32 | 0.43 |
| CCI-all vs. usual care<sup>2</sup> |  |  | -80 ± 41 | 0.05 | -81 ± 33 | 0.02 |  | -76 ± 35 | 9.9x10<sup>-7</sup> |
| **ΔLDL-C (mg·dL<sup>-1</sup>)** |  |  |  |  |  |  |  |  |  |
| CCI-all education<sup>1</sup> | 172 | 111 ± 3 | 11 (32) | 7.7x10<sup>-6</sup> | 11 ± 3 | 2.6x10<sup>-5</sup> | 113 ± 3 | 10 ± 2 | 4.9x10<sup>-5</sup> |
| Usual care<sup>1</sup> | 48 | 90 ± 4 | -11 (38) | 0.05 | -11 ± 5 | 0.03 | 90 ± 5 | -11 ± 5 | 0.02 |
| CCI-all vs. usual care<sup>2</sup> |  |  | 22 ± 6 | 0.0003 | 22 ± 6 | 0.0002 |  | 21 ± 5 | 9.9x10<sup>-5</sup> |
| **ΔHDL-C (mg·dL<sup>-1</sup>)** |  |  |  |  |  |  |  |  |  |
| CCI-all education<sup>1</sup> | 186 | 50 ± 1 | 8 (12) | <10<sup>-16</sup> | 7 ± 1 | <10<sup>-16</sup> | 50 ± 1 | 8 ± 1 | <10<sup>-16</sup> |
| Usual care<sup>1</sup> | 59 | 35 ± 2 | -2 (9) | 0.15 | -1 ± 2 | 0.69 | 37 ± 2 | -1 ± 1 | 0.41 |
| CCI-all vs. usual care<sup>2</sup> |  |  | 9 ± 1 | 1.7x10<sup>-10</sup> | 8 ± 2 | 9.9x10<sup>-6</sup> |  | 9 ± 2 | 1.2x10<sup>-8</sup> |
| **ΔTriglycerides/HDL-C ratio** |  |  |  |  |  |  |  |  |  |
| CCI-all education<sup>1</sup> | 186 | 4.3 ± 0.6 | -1.8 (9.4) | <10<sup>-16</sup> | -1.9 ± 0.9 | <10<sup>-16</sup> | 4.1 ± 0.6 | -1.6 ± 0.7 | <10<sup>-16</sup> |
| Usual care<sup>1</sup> | 59 | 12.5 ± 2.7 | 0.9 (16.1) | 0.1 | 1.2 ± 1.6 | 0.16 | 11.2 ± 2.1 | 1.0 ± 1.7 | 0.24 |
| CCI-all vs. usual care<sup>2</sup> |  |  | -2.8 ± 2.2 | 3.1x10<sup>-10</sup> | -3.1 ± 1.8 | 5.5x10<sup>-7</sup> |  | -2.6 ± 1.8 | 4.5x10<sup>-9</sup> |
| **ΔLarge VLDL-P (nmol·L<sup>-1</sup>)** |  |  |  |  |  |  |  |  |  |
| CCI-all education<sup>1</sup> | 203 | 6 ± 1 | -4 (7) | 5.6x10<sup>-15</sup> | -4 ± 1 | 1.6x10<sup>-14</sup> | 6 ± 1 | -4 ± 1 | 4.2x10<sup>-15</sup> |
| Usual care<sup>1</sup> | 68 | 12 ± 2 | 0 (8) | 0.71 | 0 ± 1 | 0.60 | 12 ± 1 | 0 ± 1 | 0.77 |
| CCI-all vs. usual care<sup>2</sup> |  |  | -3 ± 1 | 0.001 | -3 ± 1 | 0.002 |  | 3 ± 1 | 0.0007 |
| **ΔTotal LDL-P (nmol·L<sup>-1</sup>)** |  |  |  |  |  |  |  |  |  |
| CCI-all education<sup>1</sup> | 203 | 1234 ± 30 | -62 (375) | 0.02 | -57 ± 29 | 0.05 | 1235 ± 29 | -64 ± 26 | 0.02 |
| Usual care<sup>1</sup> | 68 | 1196 ± 60 | -47 (491) | 0.43 | -67 ± 53 | 0.21 | 1231 ± 57 | -57 ± 56 | 0.31 |
| CCI-all vs. usual care<sup>2</sup> |  |  | -15 ± 65 | 0.82 | 10 ± 62 | 0.87 |  | -7 ± 62 | 0.91 |
| **ΔSmall LDL-P (nmol·L<sup>-1</sup>)** |  |  |  |  |  |  |  |  |  |
| CCI-all education<sup>1</sup> | 203 | 614 ± 22 | -164 (332) | 2.2x10<sup>-12</sup> | -161 ± 24 | 4.1x10<sup>-11</sup> | 613 ± 21 | -161 ± 23 | 1.2x10<sup>-12</sup> |
| Usual care<sup>1</sup> | 68 | 724 ± 44 | 25 (370) | 0.57 | 16 ± 45 | 0.71 | 740 ± 41 | 18 ± 42 | 0.67 |
| CCI-all vs. usual care<sup>2</sup> |  |  | -189 ± 51 | 0.0002 | -177 ± 52 | 0.0007 |  | -179 ± 48 | 0.0002 |
| **ΔLDL-particle size (nm)** |  |  |  |  |  |  |  |  |  |
| CCI-all education<sup>1</sup> | 201 | 20.53 ± 0.04 | 0.23 (0.54) | 1.7x10<sup>-9</sup> | 0.23 ± 0.04 | 8.9x10<sup>-9</sup> | 20.53 ± 0.04 | 0.23 ± 0.04 | 6.0x10<sup>-10</sup> |
| Usual care<sup>1</sup> | 68 | 20.25 ± 0.07 | -0.08 (0.53) | 0.24 | -0.08 ± 0.07 | 0.25 | 20.25 ± 0.07 | -0.07 ± 0.06 | 0.25 |
| CCI-all vs. usual care<sup>2</sup> |  |  | 0.30 ± 0.07 | 4.4x10<sup>-5</sup> | 0.31 ± 0.08 | 0.0002 |  | 0.30 ± 0.07 | 3.8x10<sup>-15</sup> |
| **ΔTotal HDL-P (μmol·L<sup>-1</sup>)** |  |  |  |  |  |  |  |  |  |
| CCI-all education<sup>1</sup> | 203 | 33.2 ± 0.5 | 1.5 (4.9) | 1.2x10<sup>-5</sup> | 1.5 ± 0.4 | 2.1x10<sup>-5</sup> | 32.8 ± 0.4 | 1.5 ± 0.3 | 5.6x10<sup>-6</sup> |
| Usual care<sup>1</sup> | 68 | 29.4 ± 0.8 | -0.8 (4.7) | 0.15 | -0.8 ± 0.6 | 0.23 | 29.2 ± 0.7 | -0.7 ± 0.6 | 0.23 |
| CCI-all vs. usual care<sup>2</sup> |  |  | 2.3 ± 0.7 | 0.0004 | 2.3 ± 0.7 | 0.003 |  | 2.2 ± 0.7 | 0.0008 |
| **ΔLarge HDL-P (μmol·L<sup>-1</sup>)** |  |  |  |  |  |  |  |  |  |
| CCI-all education<sup>1</sup> | 203 | 5.3 ± 0.2 | 1.0 (2.2) | 2.5x10<sup>-11</sup> | 1.0 ± 0.2 | 4.1x10<sup>-11</sup> | 5.3 ± 0.2 | 1.0 ± 0.2 | 1.2x10<sup>-11</sup> |
| Usual care<sup>1</sup> | 68 | 3.9 ± 0.3 | 0.1 (1.6) | 0.69 | 0.2 ± 0.3 | 0.44 | 3.9 ± 0.3 | 0.1 ± 0.2 | 0.74 |
| CCI-all vs. usual care<sup>2</sup> |  |  | 0.9 ± 0.3 | 0.0002 | 0.8 ± 0.3 | 0.01 |  | 0.9 ± 0.3 | 0.0004 |
| **ΔLP-IR score** |  |  |  |  |  |  |  |  |  |
| CCI-all education<sup>1</sup> | 203 | 58 ± 2 | -14 (18) | <10<sup>-16</sup> | -14 ± 1 | <10<sup>-16</sup> | 58 ± 1 | -14 ± 1 | <10<sup>-16</sup> |
| Usual care<sup>1</sup> | 68 | 74 ± 2 | -1 (16) | 0.73 | -2 ± 2 | 0.41 | 75 ± 2 | -1 ± 2 | 0.74 |
| CCI-all vs. usual care<sup>2</sup> |  |  | -13 ± 2 | 3.8x10<sup>-9</sup> | -12 ± 3 | 6.2x10<sup>-6</sup> |  | -13 ± 2 | 6.2x10<sup>-9</sup> |
| **ΔC-reactive protein (mg·L<sup>-1</sup>)** |  |  |  |  |  |  |  |  |  |
| CCI-all education<sup>1</sup> | 193 | 5.7 ± 0.5 | -3.3 (13.4) | <10<sup>-8</sup> | -3.1 ± 1.0 | <10<sup>-16</sup> | 5.6 ± 0.6 | -3.6 ± 1.1 | <10<sup>-16</sup> |
| Usual care<sup>1</sup> | 70 | 10.4 ± 1.8 | 1.3 (13.3) | 0.94 | 0.9 ± 1.7 | 0.88 | 10.3 ± 1.6 | 1.3 ± 1.5 | 0.93 |
| CCI-all vs. usual care<sup>2</sup> |  |  | -4.7 ± 1.9 | 1.2x10<sup>-6</sup> | -4.0 ± 2.0 | 3.0x10<sup>-5</sup> |  | -4.9 ± 1.8 | 9.3x10<sup>-7</sup> |
| **ΔWBC (k•mm<sup>-3</sup>)** |  |  |  |  |  |  |  |  |  |
| CCI-all education<sup>1</sup> | 204 | 6.5 ± 0.1 | -0.7 (1.4) | 2.1x10<sup>-11</sup> | -0.7 ± 0.1 | 2.1x10<sup>-11</sup> | 6.6 ± 0.1 | -0.7 ± 0.1 | 3.2x10<sup>-11</sup> |
| Usual care<sup>1</sup> | 72 | 8.3 ± 0.3 | -0.1 (1.6) | 0.76 | -0.1 ± 0.2 | 0.74 | 8.1 ± 0.3 | -0.1 ± 0.2 | 0.76 |
| CCI-all vs. usual care<sup>2</sup> |  |  | -0.6 ± 0.2 | 0.003 | -0.6 ± 0.2 | 0.004 |  | -0.6 ± 0.2 | 0.003 |
| **Δ10-year ASCVD risk (%)** |  |  |  |  |  |  |  |  |  |
| CCI-all education<sup>1</sup> | 135 | 10.5 ± 0.7 | 1.6 (5.4) | 0.0004 | -1.5 ± 0.6 | 0.01 | 9.6 ± 0.5 | -1.3 ± 0.3 | 4.9x10<sup>-5</sup> |
| Usual care<sup>1</sup> | 55 | 12.7 ± 1.5 | 1.4 (9.3) | 0.28 | 1.1 ± 1.0 | 0.27 | 12.9 ± 1.2 | 1.2 ± 0.9 | 0.17 |
| CCI-all vs. usual care<sup>2</sup> |  |  | 03.0 ± 1.3 | 0.03 | -2.6 ± 1.2 | 0.03 |  | -2.6 ± 1.0 | 0.008 |
| **ΔCIMT-average (mm)** |  |  |  |  |  |  |  |  |  |
| CCI-all education<sup>1</sup> | 144 | 0.695 ± 0.009 | 0.002 (0.055) | 0.63 | 0.003 ± 0.004 | 0.45 | 0.685 ± 0.010 | 0.002 ± 0.004 | 0.65 |
| Usual care<sup>1</sup> | 68 | 0.680 ± 0.013 | 0.004 (0.041) | 0.37 | 0.002 ± 0.006 | 0.74 | 0.680 ± 0.013 | 0.001 ± 0.006 | 0.87 |
| CCI-all vs. usual care<sup>2</sup> |  |  | -0.002 ± 0.007 | 0.74 | 0.001 ± 0.008 | 0.87 |  | 0.001 ± 0.007 | 0.88 |
| **ΔStatin (%)** |  |  |  |  |  |  |  |  |  |
| CCI-all education<sup>1</sup> | 218 | 48.2 ± 3.4 | -3.7 (34.4) | 0.12 | -3.6 ± 2.4 | 0.13 | 46.7 ± 3.2 | -3.3 ± 2.3 | 0.15 |
| Usual care<sup>1</sup> | 73 | 64.4 ± 5.6 | 9.6 (37.9) | 0.03 | 9.5 ± 4.3 | 0.03 | 67.4 ± 5.4 | 8.8 ± 4.3 | 0.04 |
| CCI-all vs. usual care<sup>2</sup> |  |  | -13.3 ± 5.0 | 0.008 | -13.2 ± 5.0 | 0.009 |  | -12.1 ± 4.9 | 0.01 |
| **ΔAny antihypertensive medication (%)** |  |  |  |  |  |  |  |  |  |
| CCI-all education<sup>1</sup> | 218 | 56.4 ± 3.4 | -11.9<br>(42.3) | 3.2x10<sup>-5</sup> | -11.9 ± 2.9 | 3.6x10<sup>-5</sup> | 55.8 ± 3.3 | -11.4 ± 2.8 | 5.3x10<sup>-5</sup> |
| Usual care<sup>1</sup> | 73 | 60.3 ± 5.8 | 9.6 (41.4) | 0.05 | 9.6 ± 5.1 | 0.06 | 61.2 ± 5.6 | 8.3 ± 4.8 | 0.09 |
| CCI-all vs. usual care<sup>2</sup> |  |  | -21.5 ± 5.6 | 0.0002 | -21.6 ± 6.0 | 0.0004 |  | -19.7 ± 5.6 | 0.0004 |
| **ΔACE or ARB (%)** |  |  |  |  |  |  |  |  |  |
| CCI-all education<sup>1</sup> | 218 | 28.9 ± 3.1 | 0.9 (27.1) | 0.62 | 1.5 ± 1.9 | 0.42 | 30.0 ± 2.9 | 0.6 ± 1.9 | 0.76 |
| Usual care<sup>1</sup> | 73 | 21.9 ± 4.9 | 5.5 (28.3) | 0.1 | 3.7 ± 3.3 | 0.27 | 23.4 ± 4.7 | 5.0 ± 3.3 | 0.13 |
| CCI-all vs. usual care<sup>2</sup> |  |  | -4.6 ± 3.8 | 0.23 | -2.1 ± 3.9 | 0.59 |  | -4.4 ± 3.8 | 0.24 |
| **ΔDiuretics (%)** |  |  |  |  |  |  |  |  |  |
| CCI-all education<sup>1</sup> | 218 | 31.7 ± 3.2 | -9.6 (41.3) | 0.0006 | -9.5 ± 2.7 | 0.0004 | 31.3 ± 3.1 | -9.7 ± 2.7 | 0.0004 |
| Usual care<sup>1</sup> | 73 | 30.1 ± 5.4 | 5.5 (32.9) | 0.16 | 5.2 ± 4.8 | 0.28 | 33.0 ± 5.3 | 3.2 ± 4.1 | 0.44 |
| CCI-all vs. usual care<sup>2</sup> |  |  | -15.1 ± 4.8 | 0.001 | -14.7 ± 5.6 | 0.009 |  | -12.8 ± 4.9 | 0.009 |
| <sup>1</sup> Means (standard deviations) are presented. Sample sizes, means, and significance levels refer to subjects with baseline and one-year measurements for *completers*, and to 349 subjects (262 intervention and 87 usual care) for *all starters*. Significance levels for *completers* refer to one-sample t-test with or without adjustment. Untransformed triglyceride and C-reactive protein values are presented, however, their statistical significances were based on their log-transformed values. All CC refers to the CCI-web and CCI-on-site combined.<br><sup>2</sup> Mean differences ± one standard error. Significance levels refer to two-sample t-test or analysis of covariance for the differences.<br><sup>3</sup> Adjusted for sex, age, baseline BMI, baseline insulin use (user vs. non-user), and African-American race.<br><sup>4</sup> Imputed values based on 700 iterations from multivariate normal regression.<br><sup>5</sup> A significance level of P<0.0019 ensures overall simultaneous significance of P≤0.05 over the 26 variables using Bonferroni correction. |  |  |  |  |  |  |  |  |  |

**Figure 1.**
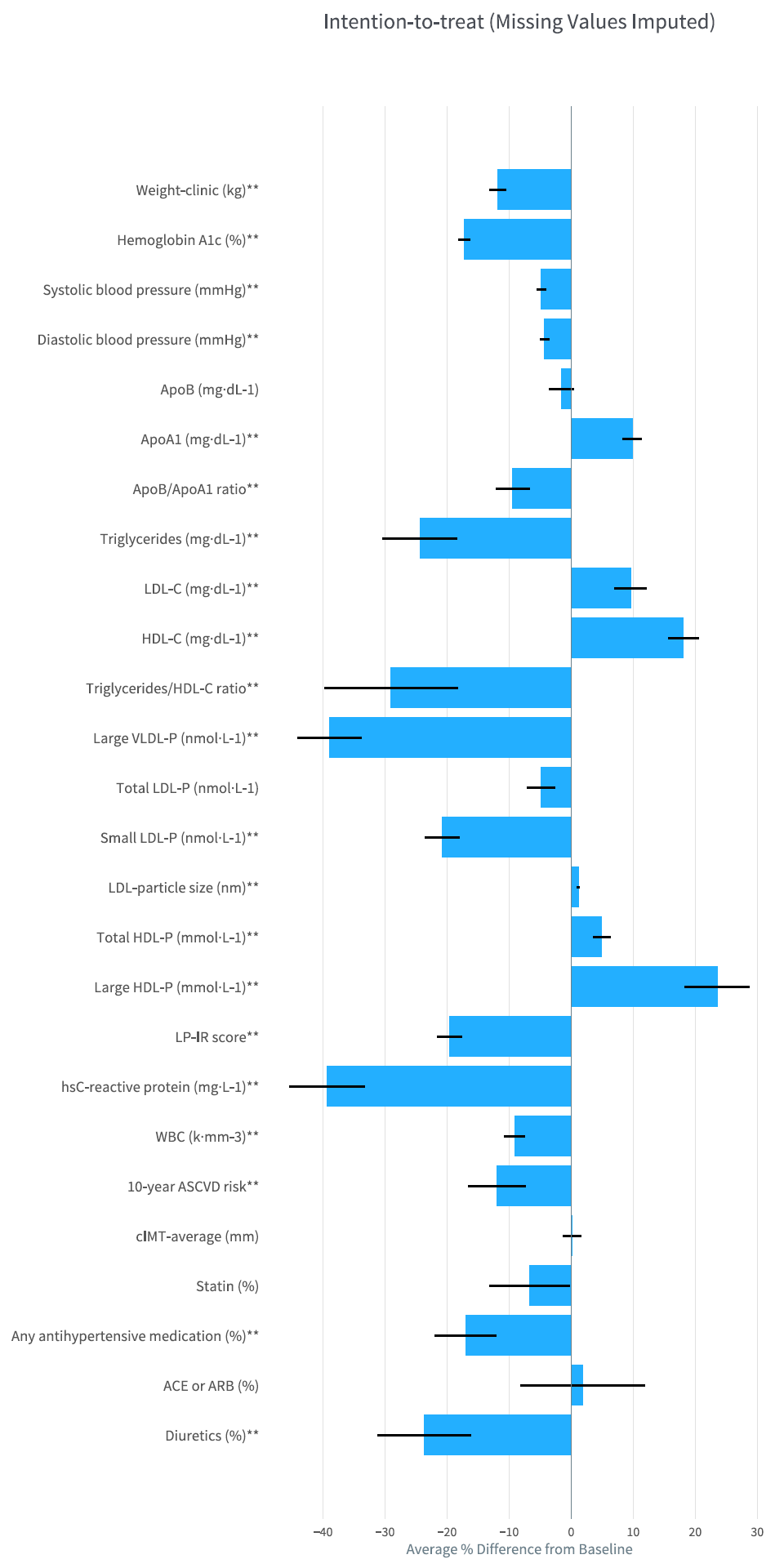
Change in biomarkers for CCI group. Bars indicate CCI group mean percent change in biomarkers based on the intention-to-treat analysis with missing values imputed. Percent change is computed as the change in mean values from baseline to one year divided by the mean baseline value. *Percent change* = 100*[(*1 year value* - *baseline value*)/(*baseline*)]. Negative values indicate a decrease from baseline to one year while positive values indicate an increase. The ** symbol after the biomarker label indicates a statistically significant within group change from baseline (P<0.0019, P adjusted for multiple comparisons). Error bars represent ± SE. *SE as Percent*= 100*[(*1 year value SE*)/(*baseline*)]

**Figure 2.**
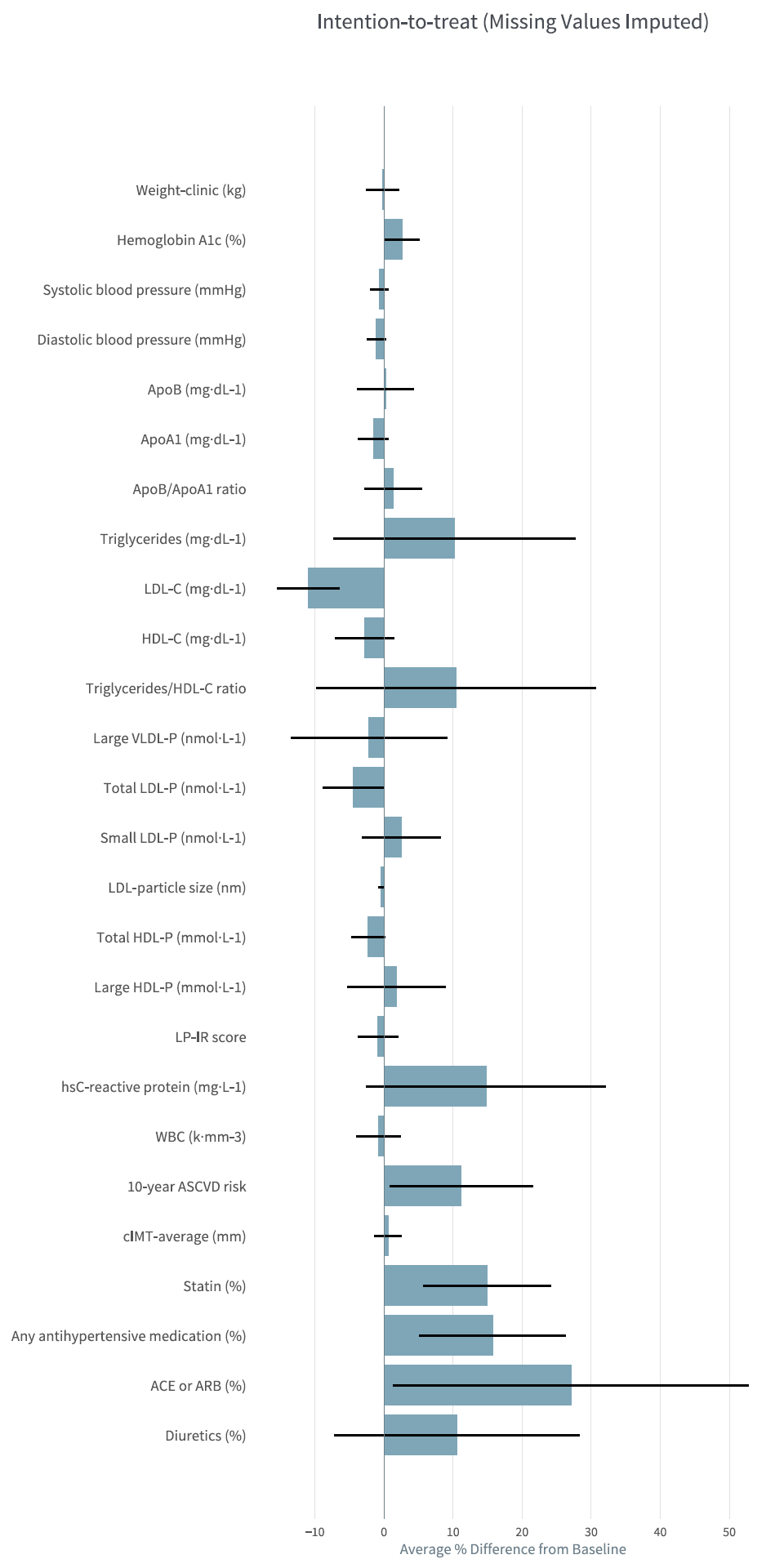
Change in biomarkers for UC group. Bars indicate UC group mean percent change in biomarkers based on the intention-to-treat analysis with missing values imputed. Percent change is computed as the change in mean values from baseline to one year divided by the mean baseline value. *Percent change* = 100*[(*1 year value* - *baseline value*)/(*baseline*)]. Negative values indicate a decrease from baseline to one year while positive values indicate an increase. (None of the within group changes were statistically significant, i.e. all P>0.0019, P adjusted for multiple comparisons.) Error bars represent ± SE. *SE as Percent*= 100*[(*1 year value SE*)/(*baseline*)]

The CCI group experienced significant reductions in systolic BP (−4.8%), diastolic BP (−4.3%), hs-CRP (−39.3%) and WBC count (−9.1%). Significant reductions were observed in overall use of antihypertensive medication (−11.4%) and diuretics (−9.6%) whereas changes in ACE or ARB (0.7%) and statin (−3.3%) use were not significant. Significant reductions were observed in both multivariate metrics: 10-year ASCVD risk (−11.9%) and LP-IR (−19.6%). There was no significant change in cIMT (averaged right and left values). In addition, changes in cIMT were not significantly correlated with baseline LDL-P or LDL-C, or changes in LDL-P or LDL-C (all p≥0.33).

One-year results from the UC group are provided in Table 2 and Figure 1. Within the UC group, after adjustment for multiple comparisons there were no significant changes at one year. After adjusting for differences in baseline characteristics and multiple comparisons, the changes observed at one year for the following biomarkers were significantly different between the CCI and UC groups (mean ΔCCI – mean ΔUC, where Δ is one year minus baseline): small LDL-P (−178 nmol·L^-1^), ApoA1 (+16 mg·dL^-1^), triglyceride/HDL-C ratio (−3.1), LDL particle size (+0.31 nm), HDL-C (+8 mg·dL^-1^), LDL-C (+22 mg·dL-^1^), hs-CRP (−4.0 mg·dL^-1^), and LP-IR (−12).

### Range of outcomes

The distribution and range of intervention response for the CCI and UC groups were compared for LDL-P, small LDL-P, large VLDL-P, ApoB, ApoA1, ApoB/ApoA1 ratio, and TG/HDL-C ratio (Supplementary Figure 1). Ranges of change observed in the CCI group were within the ranges observed in the UC group for increases in LDL-P, small LDL-P, ApoB and ApoB/ApoA1 ratio. There were two CCI participants (2/203, 1.0%) whose change in large VLDL-P exceeded the maximum observed in the UC group (15.2 nmol·L^-1^). There was one CCI participant (1/185, 0.5%) whose change in ApoA1 was less than the minimum observed in the UC group (58 mg·dL^-1^) and one CCI participant (1/186, 0.5%) whose change in triglyceride/HDL-C ratio was higher than the maximum observed in the UC group (64.9).

## Discussion

This study demonstrates that a CCI utilizing remote physician and health coach support with nutritional ketosis beneficially altered most CVD risk factors in patients with T2D at one year. Changes included: decreased small LDL-P, triglycerides, blood pressure and antihypertensive medication, hsCRP, and WBC count; increased HDL-C and LDL particle size; no change in LDL-P, ApoB, and cIMT and an increase in LDL-C. Combined with the previously reported improvements in glycemic control and reduction in obesity [10], which reduce CVD risk [49], these results demonstrate multiple additional benefits of the CCI with the exception of increased LDL-C.

Studies of dietary carbohydrate restriction, with a presumed increase in saturated fat intake, have shown modest changes in LDL-C levels [15,26-28,50]. The mean 10 mg/dL change observed in the CCI group in this study is numerically higher than the upper range of values reported by meta-analysis of lipid changes over one year related to carbohydrate restriction (−7 mg/dl to +7 mg/dl) [51]. Higher LDL-C is related to increased CVD risk [52,53], but also is *inversely* correlated with mortality in two large prospective studies and a systemic review [54-56]. Additionally, there is no evidence that increasing or decreasing LDL-C with diet interventions has any impact on mortality. LDL-C increased in the current study but both ApoB and LDL-P, measures found to be better predictors of CVD risk, did not change significantly [20-23,25,57]. In addition, the reduction in small LDL-P, increase in LDL size, and decrease in large VLDL-P that occurred in the present investigation are also associated with reduced CVD risk [58-60].

While mean response of CCI participants demonstrated an improvement in most lipid biomarkers and CVD risk factors other than LDL-C, we investigated whether a minority of participants might have unfavorable responses to the intervention. Our results suggest that a small number of participants (≤ 1%) demonstrated changes at one year outside the range of what was observed in a usual care population (Supplementary Figure 1). Thus, these results counter the concern that a significant portion of participants may have an extremely adverse reaction to the CCI (due to presumed increase in saturated fat intake) as compared to UC.

Inflammation is directly involved in all aspects of the pathogenesis of CVD [33]. High-sensitivity CRP and WBC count are widely accepted markers of inflammation and risk factors for CVD [29-32]. In addition to reducing cholesterol, reduction in inflammation may be a secondary mechanism of statins in lowering CVD risk [61-63]. The present study demonstrated a 39% reduction of hsCRP and 9% reduction in WBC count in the CCI, indicating a significant reduction in inflammation at one year. This response may be due in part to suppression of the NLRP3 inflammasome by BHB [64].

The reduction of blood pressure with concurrent reduction in antihypertensive medication was also significant. Blood pressure goals were recently reduced [65] and strong evidence exists that elevated blood pressure is a primary cardiovascular risk factor [66]. An analysis of a large T2D population suggested that antihypertensive medication may have limited effectiveness in reducing the prevalence of hypertension in these patients [67], whereas a study of weight loss interventions showed that a decrease in blood pressure predicted regression of carotid vessel wall volume [68]. Thus, additional lifestyle interventions that can augment blood pressure reduction such as the CCI described here may reduce CVD events. Additionally, the antihypertensives that were primarily decreased in the current study were shown to increase the risk for diabetes [69]. Their removal may represent further metabolic benefit.

Carotid intima media thickness (cIMT) is a non-invasive measure of subclinical atherosclerosis that is significantly associated with CVD morbidity and mortality [47,48,70,71]. However, a recent meta-analysis in 3,902 patients with T2D found that cIMT progression over an average of 3.6 years did not correlate with CVD events [70]. We found no significant change in cIMT from baseline to one year in either the CCI or UC groups. Progression or regression of cIMT may take multiple years to manifest and may require a larger cohort to achieve statistical significance [71]. In summary, the cIMT results from this study provide no evidence of vascular harm or benefit from one year of nutritional ketosis in patients with T2D.

### Strengths and limitations of the study

Prior studies have demonstrated favorable improvements in atherogenic dyslipidemia with minimal or no change in LDL-C and LDL-P following managed ketogenic diets in small short-term randomized trials. This study’s strengths include its larger cohort with high retention, prospective design and one-year duration. The study was the first to assess ApoB and ApoA1 in a T2D population adhering to a ketogenic diet. This study also has real-world application due to the outpatient setting without the use of meal replacements or food provisions.

Limitations of this study include the lack of randomization between the CCI and UC groups. In addition, the intervention provided to CCI participants was of greater intensity than UC. This was a single site study and the racial composition of study participants was predominantly Caucasian. The study was not of sufficient size and duration to determine significant differences in CVD morbidity or mortality. Since the intervention led to concurrent weight loss and improvements in cardiovascular health, it is difficult to conclude how much of the improvement can be attributed to weight loss versus other simultaneous metabolic changes. Future trials could include a longer multi-site, randomized controlled trial to allow for hard end point evaluation. Greater racial and ethnic diversity, a broader age range, and greater disease severity could also be evaluated.

## Conclusions

A T2D intervention combining technology-enabled continuous remote care with individualized plans encouraging nutritional ketosis has demonstrated diabetes status improvement while improving many CVD risk factors including atherogenic dyslipidemia, inflammation and blood pressure while decreasing use of antihypertensive mediations. Ongoing research will determine the continued safety, sustainability, and effectiveness of the intervention.

## List of abbreviations

ACE: angiotensin-converting-enzyme inhibitors
ApoA1: apolipoprotein A1
ApoB: apolipoprotein B
ARB: angiotensin II receptor blockers
ASCVD risk score: 10-year atherosclerotic cardiovascular disease risk score
BHB: beta-hydroxybutyrate
BP: blood pressure
CCI: continuous care intervention
CCI-onsite: subset of CCI participants who selected to receive onsite education
CCI-web: subset of CCI participants who selected to receive web-based education
cIMT: carotid intima media thickness
CVD: cardiovascular disease
GDR: glucose disposal rate
HOMA-IR: homeostatic model assessment of insulin resistance
hsCRP: high sensitive C-reactive protein
LP-IR: lipoprotein insulin resistance score
T2D: type 2 diabetes
UC: usual care
VLDL-P: very low-density lipoprotein particle number
WBC: white blood cell

## Declarations

### Ethics approval and consent to participate

This study was approved by the Franciscan Health Lafayette Institutional Review Board, and participants provided written informed consent.

### Consent for publication

Not Applicable

### Availability of data and material

The datasets used and/or analysed during the current study are available from the corresponding author on reasonable request.

### Authors’ contributions

SJH, ALM, WWC, JPM, SDP and JSV conceptualized and designed the study. PTW performed the formal analysis. SJH and ALM contributed to the investigation. NHB and SJH wrote the original draft. NHB and PTW created the data tables and visualizations. All authors contributed to revising and editing of the manuscript. All authors read and approved the final manuscript.

## Acknowledgements

The authors are extremely grateful to the research participants for offering their time and energy to participate in advancing scientific discovery. Thanks to Indiana University Health (IUH) staff, especially Tamara Hazbun, Monica Keyes, Danielle Wharff, Patti McKee, Joni Anderson, Zachary Roberts, Christina Selsor, and Douglas Jackson, and Virta Health staff Rachel Bolden, Sydney Rivera, and Deklin Veenhuizen for contributions to various aspects of the study including patient care, study coordination and data processing. Thanks to the health coaches who provided guidance to CCI participants: Brittanie Volk, Brent Creighton, Theresa Link, Bobbie Glon, and Marcy Abner. Thanks to Roxie McKee of IUH, Dave Gibson and Jennifer Powers of Washington University, and Teryn Sapper and staff from the Volek Laboratory at The Ohio State University for assistance in sample analysis, storage and/or transportation logistics. Thanks to Angela Fountain, Irinia Shalaurova, and Jim Otvos of LabCorp for guidance on interpreting LipoProfile results. Thank you to Ronald Krauss for guidance on lipid analysis and Ethan Weiss for critical reading of the manuscript.

---

**Supplementary Figure 1. Distribution of changes in selected biomarkers for CCI and UC completers** Histograms of changes at one year for CCI (blue) and UC (gray) are overlaid. Very few (≤1%) CCI participants demonstrated changes in an undesirable direction at one year that was outside the range of changes observed in the UC group for key lipid and lipoprotein particles. (A) Apolipoprotein B (B) Apolipoprotein A1 (C) Apolipoprotein B/Apolipoprotein A1 ratio (D) LDL-P (E) Small LDL-P (F) Large VLDL-P (G) Triglyceride/HDL-C ratio

